# Defining the data gap: what do we know about environmental exposure, hazards and risks of pharmaceuticals in the European aquatic environment?

**DOI:** 10.1101/2023.07.10.548305

**Authors:** F.D. Spilsbury, P.A. Inostroza, P. Svedberg, C. Cannata, A.M.J. Ragas, T. Backhaus

## Abstract

Active pharmaceutical ingredients (APIs) and their transformation products inevitably enter waterways where they might cause adverse effects to aquatic organisms. Identifying the potential risks of APIs in the environment is therefore a goal and current strategic direction of environmental management described in the EU Strategic Approach to Pharmaceuticals in the Environment and the Green Deal. This is challenged by a paucity of monitoring and ecotoxicity data to adequately describe risks.

In this study we analyze measured environmental concentrations (MECs) of APIs from 5933 sites in 25 European countries as documented in the EMPODAT database or collected by the German Environment Agency for the time period between 1997 to 2020. These data were compared with empirical data on the ecotoxicity of APIs from the U.S. EPA ECOTOX database. Although 1763 uniquely identifiable APIs are registered with the European Medicines Agency (EMA) for sale in the European Economic Area (EEA), only 312 (17.7%) of these are included in publicly available monitoring data, and only 36 (1.8%) compounds have sufficient ecotoxicological data to perform an EMA-compliant ERA. Among the 27 compounds with sufficient exposure and hazard data to conduct a single substance risk assessment according to EMA guidelines, four compounds (14.8%) had a median risk quotient (RQ) > 1. Endocrine disruptors had the highest median RQ, with 7.0 and 5.6 for 17α-ethinyl-estradiol and 17β-estradiol respectively.

A comparison of *in-silico* and empirical data for 72 APIs demonstrated the high protectiveness of the current EMA guidelines, with predicted environmental concentrations (PECs) exceeding median MECs in 98.6% of cases, with a 100-fold median increase.

This study describes the data shortfalls hindering an accurate assessment of the risk posed to European waterways by APIs, and identifies 68 APIs for prioritized inclusion in monitoring programs, and 66 APIs requiring ecotoxicity testing to fill current data gaps.

**Highlights:**

- 1763 medicines are EMA-approved for sale in the EEA
- The data gap is 1201 APIs (68%) that have no ecotoxicity or public monitoring data
- Only 27 APIs (1.5%) have sufficient empirical data for risk assessment.
- ERA using 23 years of EU monitoring data shows four compounds with a median RQ > 1
- Data gap APIs prioritized for monitoring programs (68) and ecotoxicity testing (66)

## 1. Introduction

Water quality is a “heritage which must be protected” *EU Directive 2000/60/EC, Water Framework Directive). Reduction of the risk posed by chemicals in the environment has seen much policy-driven progress, including the zero pollution ambition of the Green Deal (EU Commission, 2019a). The EU Strategic Approach to Pharmaceuticals in the Environment (EU Commission, 2019b) aims to conduct an “investigation to address the potential risks from pharmaceutical residues in the environment”, and to “identify remaining knowledge gaps, and present possible solutions for filling them”. The EU Chemicals Strategy for Sustainability (EU Commission, 2020) aims to “develop a framework of indicators to monitor the drivers and impacts of chemical pollution”, and prioritizes chemicals such as pharmaceuticals that “affect the reproductive or the endocrine system, or are persistent and bio-accumulative”.

Estimations of the total number of active pharmaceutical ingredients (APIs) in global use range from 3,500 to 10,000 (Boxall *et al*., 2012, Dong *et al*., 2013; Hughes *et al*., 2013; Caldwell *et al*., 2014; Mansour *et al*., 2016). Within the European Economic Area (EEA), approximately 2000 APIs are approved for sale (Perrazolo *et al*., 2010). The number of APIs that are causing environmental risk, however, is considerably lower, partly because consumption volumes differ vastly. For example, Kümmerer estimated that 90% of the consumption volume in Germany can be attributed to only 900 APIs (Kummerer, 2008).

APIs used for treating human disease are ubiquitous in waterways (Pereira *et al*., 2020a), entering mainly via effluent from waste water treatment plants (WWTP) (Schwarz *et al*., 2021). Efficacy of API removal by WWTP depends on the technology available (Hollender *et al*., 2009), and studies of WWTP effluent discharging to European waterways report detectable concentrations of carbamazepine, ibuprofen, diclofenac, caffeine (Loos *et al*. 2009, 2013) and antibiotics (Felis *et al*., 2020). The concentration of lipophilic APIs in surface waters is attenuated by sorption to particulate matter, whereas levels of hydrophilic compounds are moderated by dilution (Pereira *et al*., 2020b), largely dependent on river water flow rates (Burns *et al*., 2017).The number of APIs typically detected in surface waters ranges from 200 to 600 (Hughes et al., 2013; Kuster and Adler, 2014), and a comprehensive review of global environmental monitoring data by the German environmental agency (UBA) found 771 APIs present at concentrations above their respective analytical limits of reporting in 75 countries across 5 continents (aus der Beek et al., 2016; Dusi et al., 2019).

Non-detection of an API does not necessarily mean that it is actually absent. Instead, the limits of detection (LOD) of the employed analytical methods may be insufficient to detect an API at a concentration that might cause environmental concerns. Current guidelines recommend an LOD 30% below environmental quality standards (Coquery *et al*., 2005; EC 2009; ISO 13530:2009). However, for many chemicals included in European monitoring programs, analytical techniques are insufficiently sensitive (Brack *et al*., 2017).

The guidelines of the European Medicines Agency (EMA) for environmental risk assessments (ERA) of APIs (EMA, 2018) are built on a two-tier structure comprised of a low data-demand initial screen (Phase 1) which uses modelled data to determine if an API is likely to be sufficiently bioaccumulative (Log K_OW_ > 4.5) or if the API is predicted to be present in the environment at concentrations exceeding 0.01 µg/L. A more comprehensive ERA (Phase 2) with inherently higher data demands is implemented if either of those conditions is fulfilled. Antibiotics and hormones always require a Phase 2 ERA. As a minimum, chronic no observed effect concentration (NOEC) or chronic EC10 values must be available for three trophic levels (algae, daphnids and fish) in order to calculate a Predicted No Effect Concentration (PNEC) as an estimate of the environmental hazard of an API.

There is a paucity of adequate empirical ecotoxicological data available for the majority of APIs found in European water bodies (Backhaus, 2016), hampering large-scale environmental risk assessments and prioritizations.

Similarly, exposure data are also lacking, with inconsistent or incomplete monitoring data available for the majority of the almost 2000 APIs approved for sale in the EEA (Dulio et al., 2018; Brack et al., 2017, 2019). In the absence of analytical monitoring data, predicted environmental concentrations (PEC) can be estimated by a variety of models, with varying data demands and varying reliability (Burns et al., 2017, 2018). In-line with the precautionary principle, a worst-case scenario model based on maximum recommended pharmaceutical daily doses is prescribed by the EMA for deriving PECs (EMA, 2018).

In this study we provide a large-scale overview of the knowledge on the environmental risks of APIs in European surface waters. In particular, we analyze the data gaps that hamper large-scale ERA and API prioritization. In order to keep the study and its conclusions transparent and available for follow-up studies when new data become available, the analysis is based entirely on publicly available data, curated from the European Medicines Agency (EMA), U.S. EPA ECOTOX database, NORMAN-EMPODAT (EMPODAT), German Umweltbundesamt (UBA) and the World Health Organization (WHO). All data analysis scripts are made available at GitHub (https://github.com/collections/open-data/TO_BE_UPLOADED ON ACCEPTANCE). We also provide worst-case PECs according to EMA’s Phase 1 screening, in order to assess whether EMA’s approach is sufficiently conservative, in comparison to empirical monitoring data.

## 2. Materials and Methods

All analyses were performed using R statistical software (version 4.2.1). The final data analysis scripts are available at https://github.com/collections/open-data/TO_BE_UPLOADED ON ACCEPTANCE.

### 2.1 Data Compilation

#### Pharmaceuticals Registered for Sale in the EEA

A list of all pharmaceuticals centrally registered at EMA for sale in the EEA was compiled from the publicly available list of marketing authorization holders, known as the Article 57 database, obtained from the EMA website (EMA, 2023). The Article 57 list contains producer-submitted information, and includes product names, the active pharmaceutical ingredients, administration routes and contact details of the company authorized to produce and market the medicine. Identifiers for each of the pharmaceuticals such as the Chemical Abstracts Service (CAS) number, International Chemical Identifier (InChI) and its hashed InChIKey counterpart, and Simplified Molecular Input Line Entry System (SMILES) were obtained via name-match searches of the PubChem (PubChem, 2022) or Drugbank (Drugbank, 2022) websites. Treatment categories for the pharmaceuticals included in this study were allocated using Anatomical Therapeutic Codes (ATC) obtained from the WHO (WHO, 2022). ATCs are a clinical use categorization system with four tiers. Some compounds may have several ATCs, and in these cases, the primary categorization described in literature was used. All herbal medicines (EMA, 2022b) and dietary supplements were excluded from the study, as were any compounds without a CAS number and therefore unable to be uniquely identified.

### 2.2 Ecotoxicological Data

Ecotoxicological data were retrieved from the U.S EPA ECOTOX database (EPA, 2022). To ensure capture of other ecotoxicological data not included in ECOTOX, a Web of Science (WoS, 2022) literature search was performed for a selection of APIs reported in monitoring data but lacking any ecotoxicity data using the following search string: *“TS=(Name) AND TS=(EC10* OR EC05* OR NOEC* OR LOEC* OR MATC OR TLm OR ChV OR ECx OR bioassay*)”*, and checking the title and abstract of returned results. Finally, we monitored the availability of regulatory ecotoxicity data in the Registration, Evaluation, Authorization and Restriction of Chemicals (REACH) database and the European Public Assessment Reports (EPARs) published by the EMA.

Data were curated for quality by omitting records which lacked exposure durations, measurement endpoints, appropriate units or a reference for the source of the data. Only data which met the intention of the EMA guideline (EMA, 2018) for the derivation of predicted no effect concentrations (PNECs) were included. Briefly, chronic experimental toxicity data must be available for NOEC or EC10 endpoints describing adverse impacts on the reproductive ability necessary to maintain a viable population for three trophic levels: algae, daphnia and fish. Acceptable toxicity data for algal species included test durations exceeding 3 days with measured endpoints for biomass/abundance, growth or growth rate inhibition (OECD 201). Where available, data from the four algal species described in OECD 201 were used: *Raphidocelis subcapitata* (also known as *Pseudokirchneriella subcapitata*), *Desmodesmus subspicatus, Navicula pelliculosa* and *Anabaena flosaquae*. However, where appropriate toxicity data was lacking, ecotoxicity data for other species of algae were used, namely *Chlamydomonas reinhardtii* for the antidepressant phenobarbital. For daphnia, data with observations of reproductive success including hatching rates, progeny counts, survival and length with exposures exceeding 6 days were used (OECD 211). Where appropriate data for *Daphnia magna* were unavailable, other comparable species of crustaceans such as *Ceriodaphnia dubia* (waterflea), *Hyalella azteca* (scud) or *Gammarus locusta* (scud) were accepted. For fish, data with a minimum 7-day exposure for fish larvae (OECD 210) with measurements of hatch rates, survival, progeny counts, length and mortality for the following species commonly used in ecotoxicity testing were used: *Danio rerio* (zebrafish), *Pimephales promelas* (fathead minnow), *Oryzias latipes* (Japanese medaka) or *Rutilus* (common roach).

### 2.3. Environmental Monitoring Data

Raw data from the environmental monitoring of European surface waters were amalgamated from two publicly available data compilations: the EMPODAT (NORMAN) database (EMPODAT, 2022) and the UBA Measured Environmental Concentrations database (UBA, 2022). Data were available spanning a 23-year period from 1997 to 2020. Data which lacked sampling date, country, or CAS numbers were excluded from the analysis. Only monitoring data from countries which are part of the EEA were included in the study, notably excluding the United Kingdom. Duplicate records (e.g. those in UBA which also appear in the EMPODAT database) were also removed using the source field in the EMPODAT database. Only pharmaceuticals uniquely identifiable by CAS number, and authorized for sale in the EEA by the EMA and listed in the Article 57 database were included in the study. Metals (e.g. silver, copper, calcium, magnesium, zinc and iron) and their salts were excluded from the analyses, as were dietary supplements and vitamins.

### 2.4. Risk Assessment

Only empirical data were used for the assessment. Risk was calculated using measured environmental concentrations (MEC) as the exposure component, and for the hazard component PNECs were calculated by applying an assessment factor (AF) of 10 to the lowest chronic NOEC or EC10 value for the most sensitive species from three trophic levels: algae, daphnia and fish (Eq.1), compliant with EMA guidelines for environmental risk assessment (EMA, 2018).

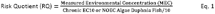

### 2.5. Prioritizing the Unknowns

APIs from EMA’s Article 57 list that are approved for sale in the EEA but were not included in the environmental monitoring data were prioritized using the Phase 1 requirements of the EMA guidelines (EMA, 2018).

Experimentally derived LogK_OW_ values obtained from PubChem (PubChem, 2022) were used in preference to estimated values to evaluate bioaccumulation potential of the APIs. The “worst-case” scenario model specified in the EMA guidelines was used to estimate environmental concentrations (Eq. 2), using defined daily dose (DDD) information obtained from the WHO (WHO, 2022). Clinical use categorizations were derived from ATC codes, also obtained from the WHO (WHO, 2022). Default values used for penetration factor (proportion of the population using an API), dilution factor (dependent on river flow rates) and the amount of daily wastewater produced by the population were 1%, 10 and 200 L/inh/day, respectively (EMA, 2018).

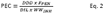

Where: DDD is the defined daily dose (maximum recommended adult dose)

F_PEN_ is the penetration factor (by default 1% of the population)

DIL is the dilution factor (by default 10)

WW_INH_ is the daily wastewater produced per inhabitant (by default 200 L/inh/day)

Compounds from the Article 57 list which were detected in European rivers, but which lack sufficient toxicity data to perform a risk assessment were prioritized by the number of detections reported in monitoring data from 1997 to 2020, and detection rate in European waterways, and the number of trophic levels (algae, daphnia or fish) for which relevant chronic NOEC or EC10 data are unavailable.

### 2.6. Comparison of *in-silico* and empirical exposures

The protectiveness of the EMA model for PEC calculation and its 0.01µg/L threshold for Phase 1 screening was investigated by comparing modelled PEC estimations of exposure against median MECs for the whole of the EEA. Individual compounds were included on the basis that a DDD was available and therefore enabling the calculation of a PEC (Eq. 2), and they were reported in concentrations exceeding their respective LOD in EMPODAT and UBA monitoring data. The protectiveness of the PEC for individual APIs was quantified as the ratio of PEC/median MEC.

## 3. Results and Discussion

### 3.1. Data-gap characterization

We identified a total of 1763 non-herbal pharmaceutical products by CAS number from the Article 57 list of medicines authorized for sale in the EEA. These APIs were from 11 different treatment categories as described by their respective ATC codes (Figure 1). Some APIs were included in more than one treatment category.

**Figure 1:**
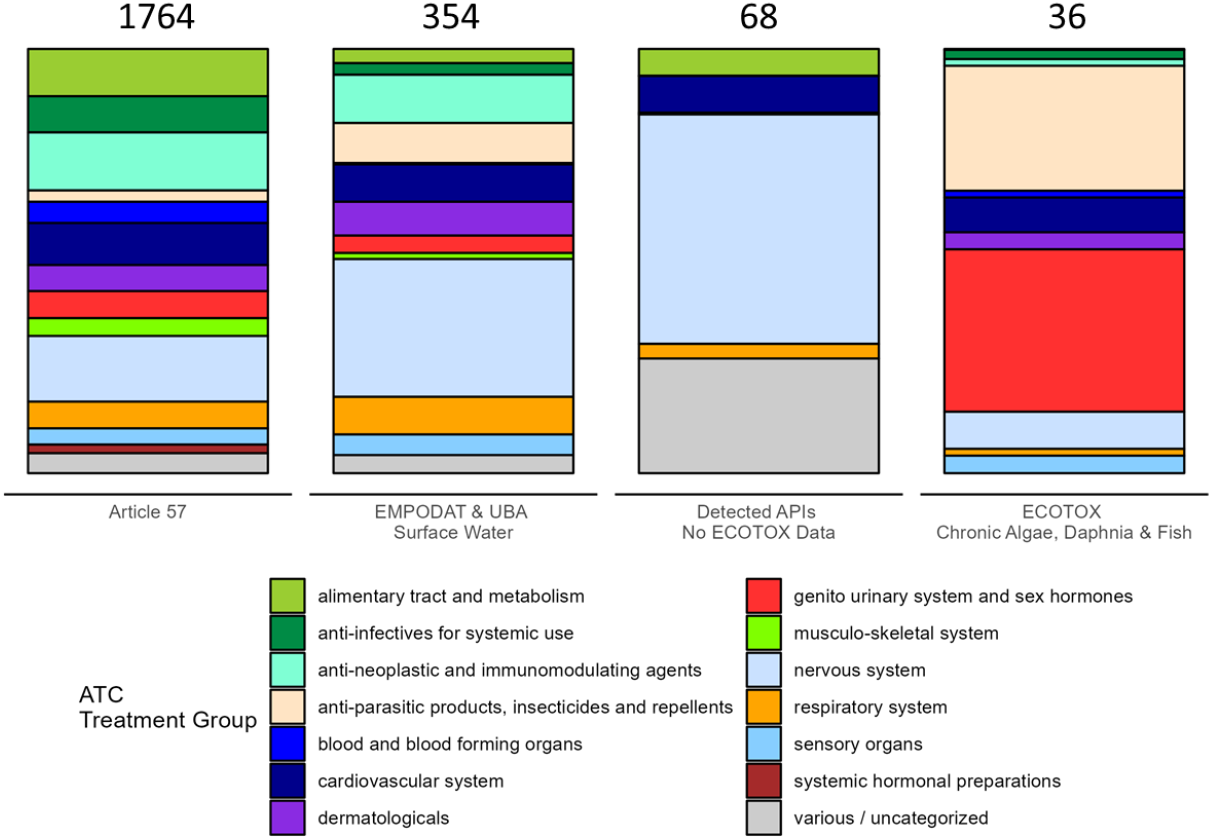
Treatment groups (ATC groups) and number of APIs for four categories of data availability: APIs listed in the EMA Article 57 list, APIs included in monitoring studies, APIs detected in monitoring studies but lacking ecotoxicity data, and APIs for which chronic ecotoxicity data spanning 3 trophic levels (algae, daphnids and fish) are available in ECOTOX and other literature.

Many pharmaceuticals and/or their degradation products are not removed from effluents during sewage treatment plant processes (Hollender *et al*, 2009; Loos *et al*, 2009, 2013; Felis *et al*, 2020), and are therefore be expected to occur in surface waters, particularly in highly urbanized areas. Of the 237,939 EMPODAT and UBA records (1997 to 2020), a total of 111492 (47%) entries report above the corresponding limits of detection (LOD) (Figure 2), similar to detection rates reported in waste water treatment effluent (Loos *et al*., 2013). Sampling intensity varies between countries (Figure 2), and between years, likely a result from different levels of monitoring activities as well as different data collection efforts (submission to EMPODAT and/or publication in the open literature so that it can be incorporated into the UBA database).

**Figure 2:**
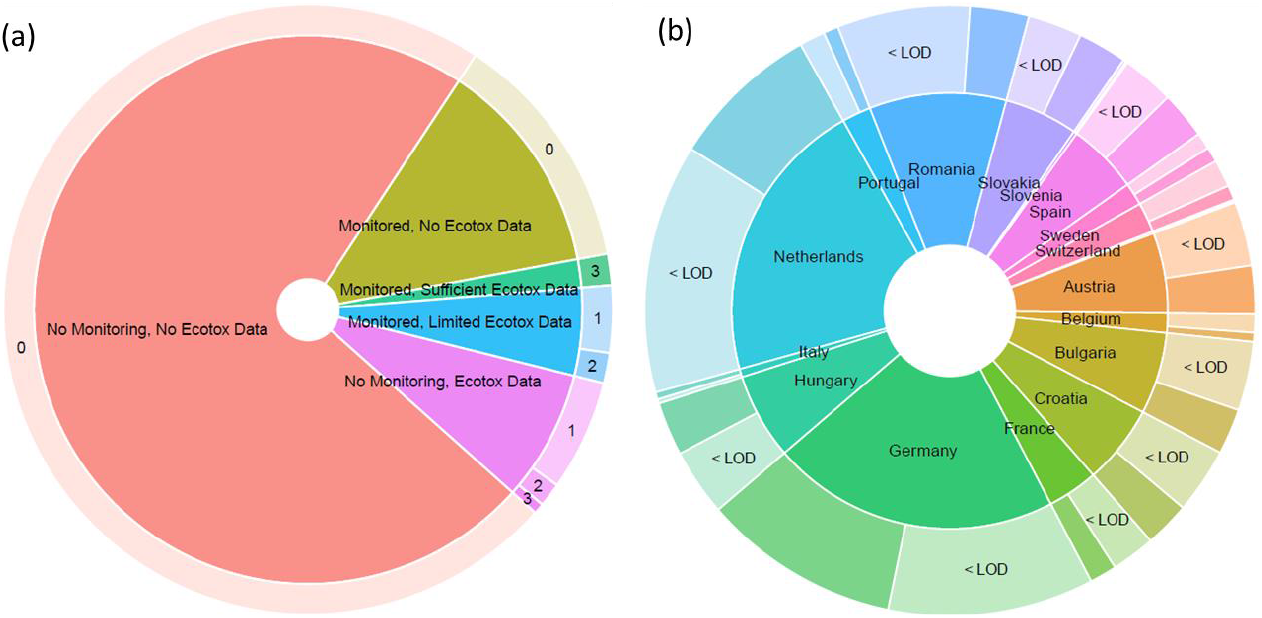
Overview of monitoring and ecotoxicity data for 1,764 Article 57 pharmaceuticals in the EEA. (a) monitoring (EMPODAT & UBA) and ecotoxicological (ECOTOX) data availability. Outer ring describes the number of species groups (algae, daphnids, fish) included in the available ecotoxicological data. (b) Publicly available monitoring data by country, and overall detection rates.

Of the 1763 APIs listed on the EMAs Article 57 list, 1201 (68% of the total) are neither listed in the EMPODAT and UBA databases, nor have any ecotoxicity data available from the EPA ECOTOX database (Figure 1). A total of 354 APIs (20% of the total) were monitored at some point in the 23-year span that is covered by the data in the EMPODAT and UBA databases, with 68 (3.9% of the total) reported as being present at least once above their respective LOD (Figures 1,2). Of these, only 36 APIs (2% of the total) meet the minimum data requirements specified in the EMA guidelines to calculate a PNEC (EMA, 2018) which requires chronic ecotoxicity data for algae, daphnia and fish (Figures 1 & 2). Interestingly, although APIs used to treat disorders of the nervous system (e.g. antidepressants and opioids) only comprise 15% of the APIs on the Article 57 list, 31% of the EU surface water monitoring data relate to these compounds, probably as a consequence of the specific targeting of pre-selected chemicals of concern by monitoring regimes (Dulio *et al*. 2016; Dulio *et al*., 2018). The 68 compounds reported to be present in the environment, but for which no ecotoxicity data were found are also dominated by APIs whose ATC code indicates an interaction with the nervous system. Conversely, the APIs for which chronic ecotoxicological data are available for all three taxonomic groups, as required for an EMA-compliant ERA are dominated by endocrine disruptors (37%) and the group of parasiticides, insecticides and repellants (28%) (Figure 1).

Insufficient ecotoxicological data are available for 52 of the 354 APIs that are found in detectable concentrations in European surface waters on more than 50 occasions between 1997 and 2020 (Table S1). Of these, we could find neither acute nor chronic toxicity data for 24 APIs. A Web of Science literature search yielded no additional ecotoxicity data not already included in ECOTOX. The anti-epileptics gabapentin and lamotrigine were detected in European waterways on 2884 and 992 occasions respectively, with detection rates exceeding 95%. Valsartan, a medication for the treatment of high blood pressure, was recorded above its LOD 1230 times with a detection rate of 91.1%. Atropine, desmethyldiazepam, dihydrocodeine, acebutolol, sitagliptin and celiprolol also had detection rates exceeding 90%.

### 3.2. Risks to the Aquatic Environment

A total of 27 of APIs included in the EMA’s Article 57 list meet the minimum data requirements to allow the estimation of risk to the aquatic environment, i.e. sufficient chronic ecotoxicological data and reported environmental concentrations. Risk quotients (RQs) were calculated for 66,393 reported concentrations of individual APIs from 22 countries, with 21,720 (32.7%) exceeding the risk threshold of RQ =1 (Figure 3). PNECs used in risk calculations and the underlying ecotoxicity data used to derive them are presented in Table S1, and are generally similar to PNEC values reported in other studies (e.g. Loos *et al*., 2018).

**Figure 3:**
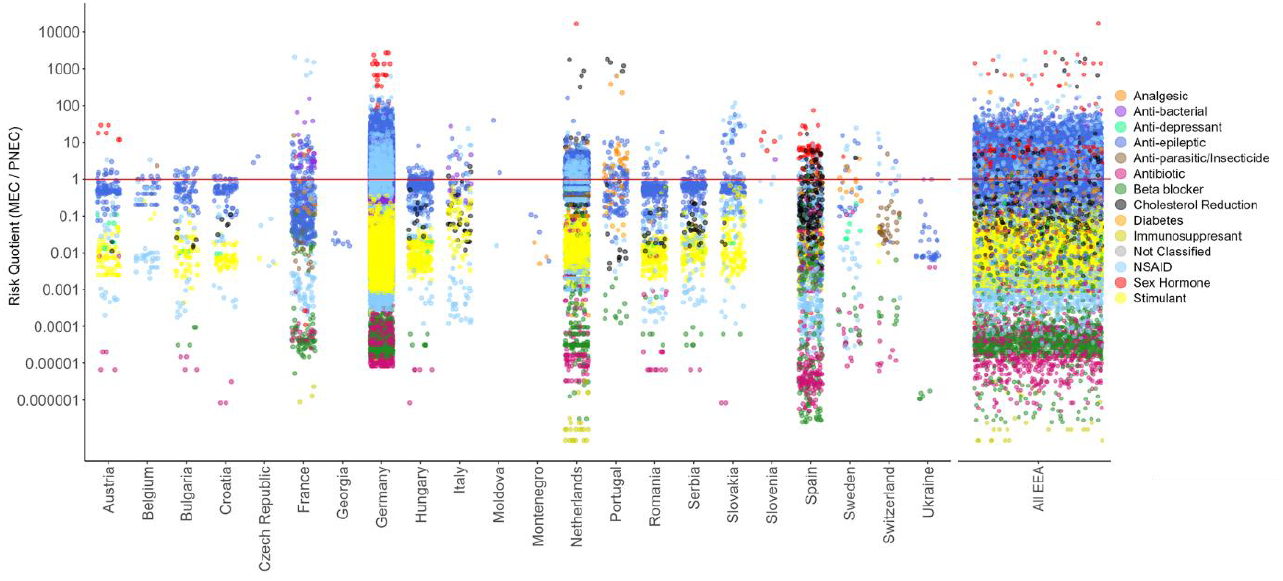
Risk assessment of 27 pharmaceuticals detected in European surface waters between 1997 and 2020. The red line is RQ = 1.

Some general trends were consistent between countries: caffeine and aspirin were reported in most countries, but generally at concentrations clearly below the PNEC (median RQs 0.006 and 0.004 respectively). Although caffeine is listed as a pharmaceutical by the EMA, the majority of concentrations found in European surface waters likely originate from the consumption of caffeine-containing beverages. Anti-depressants and non-steroidal anti-inflammatory drugs (NSAIDs) were reported in 18 of 21 countries at concentration levels often approaching or even exceeding their respective PNECs. The highest RQ recorded for any analyte was in the Netherlands in 2011, where 17α-ethinyl-estradiol was recorded at a staggering 16,666 times the threshold concentration. This is in agreement with other studies, which also report very high risk from sex hormones (Pereira *et al*., 2020b). Hormones are by their nature compounds which cause a large effect at very small doses (Runnalls *et al*., 2010).

Of the 27 compounds with sufficient ecotoxicity data, four (14.8%) have a median RQ > 1 (Figure 4). This is slightly higher than a previous study (Kuster and Adler 2014), which reported that around 10% of APIs are present in European waterways at levels of environmental concern. The two compounds with the highest median RQ were the sex hormones 17α-ethinyl-estradiol and 17β-estradiol.

**Figure 4:**
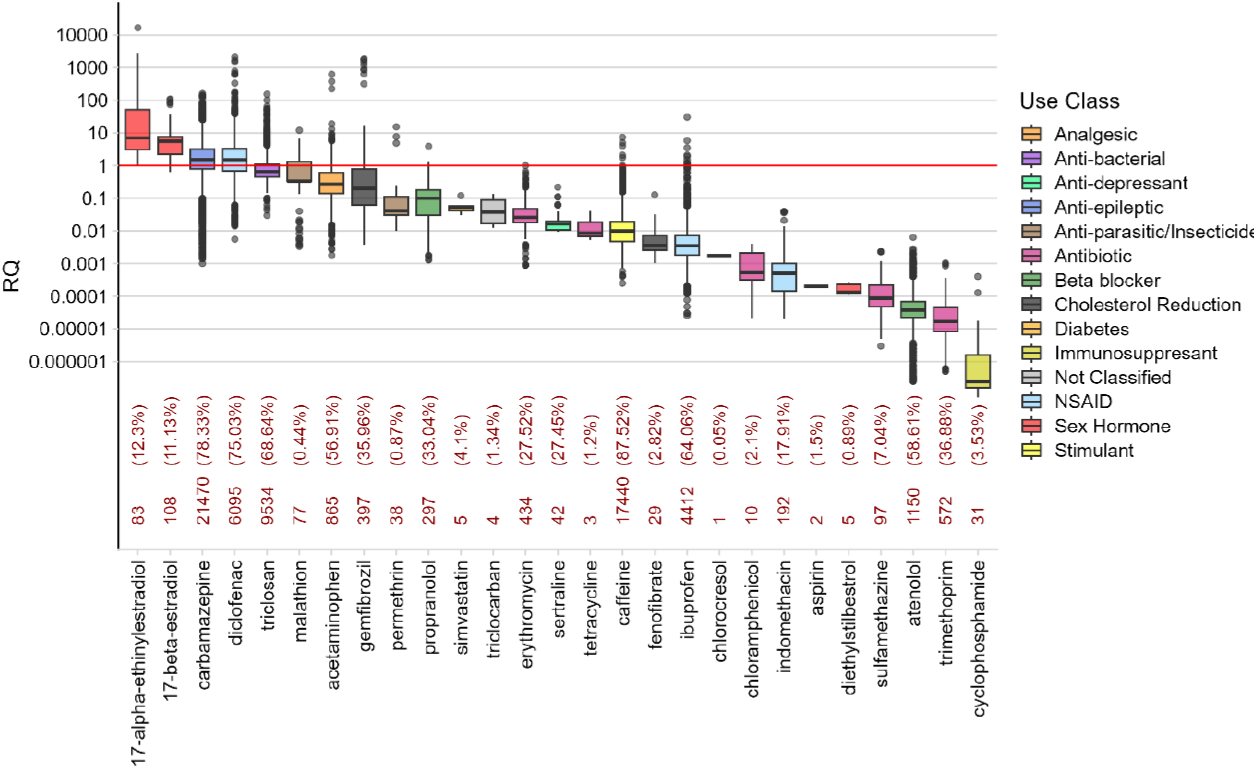
Risk of 27 pharmaceuticals detected in European rivers, ranked by median RQ. Red line indicates the risk threshold (RQ = 1). Numbers in red are number of detections (1997 – 2020), and their respective detection rate.

However, these two compounds were detected in only 12.2% and 11.1% of samples respectively. Detection frequencies range from 0.05% (chlorocresol, an anti-fungal agent) to 87.5% (caffeine) (Figure 4). Of particular note, detection rates for diclofenac and erythromycin were 75.0% and 27.5% respectively, similar to those reported in WWTP effluent studies (Loos *et al*., 2018), while 17α-ethinyl-estradiol detection rates were lower (Loos *et al*., 2018). The anti-epileptic drug carbamazepine and the NSAID diclofenac have a median RQ > 1, with 21,470 and 6,095 detections between 1997 and 2020 respectively, and high detection rates exceeding 75% (Figure 4).

The exposure of some of the APIs included in the risk analysis are likely not solely attributable to medical use. Malathion and permethrin, although the active ingredient in treatments for headlice or parasitic infections, have also been used as agricultural pesticides before their use was discontinued in Europe in 2003. Also, 17β-estradiol is a naturally occurring hormone, and its detection in surface waters is therefore not exclusively attributable to pharmaceutical use.

Previous studies have used in-silico methods such as quantitative structure-activity relationship (QSAR) modelling to fill the data-gap (Posthuma at al., 2019) with varying degrees of reliability. Another recent study seeking to define the data gap for pharmaceuticals in European rivers have used confidential toxicity data sourced from industry (Schwarz et al., 2021), making verification impossible. A risk assessment by Davey et al. (2022) of 702 APIs categorized as psychopharmaceuticals by their anatomical therapeutic code (ATC) used a combination of probabilistic and deterministic methods such as species sensitivity distributions, and the application of acute-to-chronic conversion factors to maximize the amount of available ecotoxicity data, enabling the risk to be estimated for 72 psychopharmaceuticals and illicit drugs detected in European rivers.

However, the use of conversion factors to provide ecotoxicity values equivalent to the EMA-mandated chronic NOEC or EC10 may introduce a source of error. Empirical data, while in relatively limited supply, remains the gold standard for ERA.

### 3.3. Comparison of modelled and measured exposures

The data demands of the EMA model for estimating PECs (maximum recommended daily dose (WHO, 2023); see Eq. 2) were met by 72 APIs, allowing comparison with the MECs reported in EMPODAT and UBA databases. For 71 (98.6%) APIs, PECs exceeded their respective median MEC (Figure 5). The median ratio of PEC/MEC was 100 (Figure S2), demonstrating that PECs are conservative across different API types and treatment uses. In line with the precautionary principle, 70 APIs (97.2%) exceed the 0.01 µg/L PEC threshold for Phase 1 EMA risk assessment, compared to only 27 (37.5%) if the exposure assessment is based in empirical data. Forty-five APIs exceed the EMA ERA Phase 1 exposure threshold by the EMA-mandated model, which would otherwise not require the data-intensive rigors of a comprehensive Phase 2 risk assessment under the EMA guidelines when considering their environmental exposure by empirical means.

**Figure 5:**
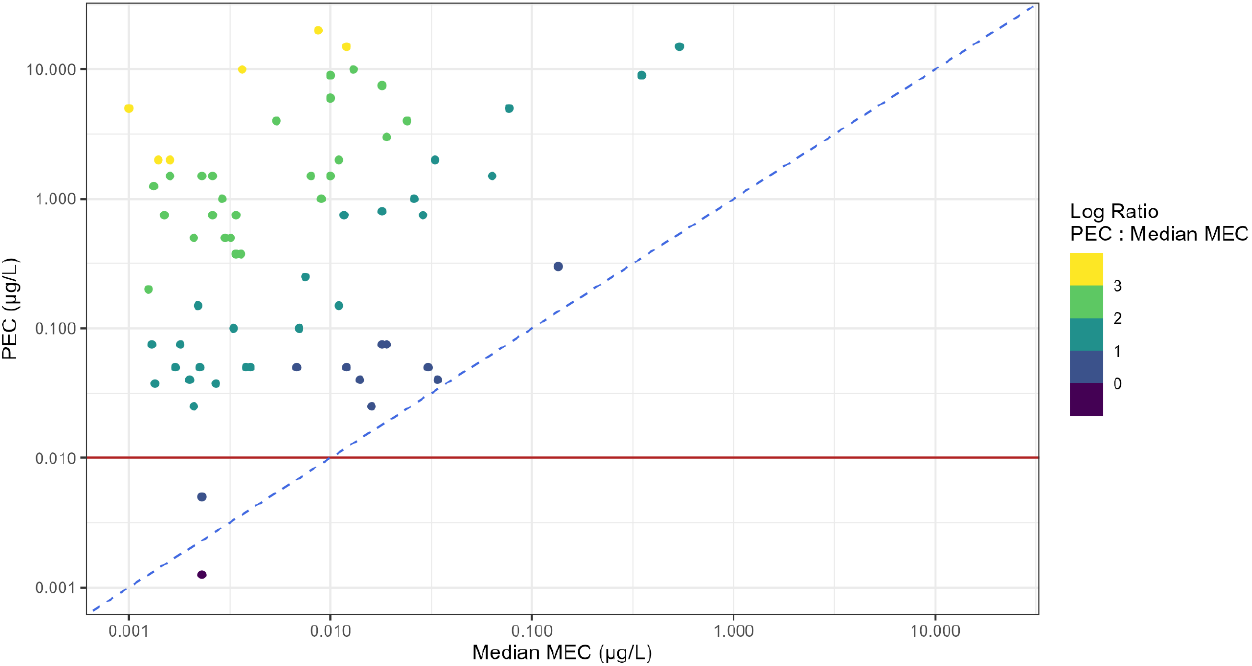
Comparison of modelled predicted environmental concentration (PEC) with median measured environmental concentration (MEC) (1997 – 2020). Blue dotted line shows the 1:1 ratio. Red line is the EMA threshold for Phase 1 risk assessment (0.01 µg/L)

The DDD data from which we derived PECs are available for 2649 APIs, 440 of which are listed in the EMA’s Article 57 list but lack empirical exposure data, and are therefore candidates for *in-silico* PEC estimation. Unsurprisingly, 431 APIs (98.0%) exceed the 0.01 µg/L EMA ERA Phase 1 threshold (Table S3). Other APIs lacking empirical data failed the EMA ERA Phase 1 screen due to a lipophilicity (LogK_OW_ > 4.5) indicating their potential to bioaccumulate (17.1%), or were classified by their ATC as being hormones (5.5%) or antibiotics (8.5%).

The EMA guidelines for ERA use worst-case scenario PEC estimations in its Phase 1 screening and is an overestimation of the concentrations of APIs likely present in surface waters of the EEA. A number of studies have proposed improvements to the EMA guidelines, such as considering transformation products (Escher and Fenner 2011) and mixture effects (Backhaus *et al*., 2014, 2016; Agerstrand *et al*., 2015). QSAR-derived data for the ecotoxicity of APIs may underestimate the ecotoxicity of APIs by a factor of ten (Sanderson *et al*, 2007; Kar *et al*, 2018), implying that high assessment factors are appropriate when estimating ecotoxicity for ERA purposes using QSAR values to fill data gaps (Esher *et al*, 2011). Even detailed exposure models with high data demands that consider local usage rates, metabolism, WWTP elimination rates and dilution rates based on seasonal river flow underestimate the concentration of pharmaceuticals in the environment (Burns *et al*., 2017, 2018). While such improvements may reduce the number of compounds proceeding to Phase 2 ERA under the current EMA guidelines, the empirical data demands to develop such models for exposure and ecotoxicity are high, and can be regarded as another data-gap requiring a bridge.

### 3.4. Prioritizing APIs for further action

APIs should be prioritized for further action, depending on the specific data gaps. 68 APIs are detected in European freshwater ecosystems, sometimes frequently, but their ecotoxicological hazard profiles are unknown (Table S2). Those compounds should therefore be prioritized for future ecotoxicological studies.

We identified 65 APIs with a problematic hazard profile and a use pattern that should be prioritized for future monitoring studies (Table S3). These are APIs with a PEC > 0.01 µg/L that are either used for their hormonal action, for their biocidal mode of action (insecticides, antimycotics, insect repellents, etc) or for which other target genes are conserved in environmental organisms (Gunnarsson et al 2019).

## 4. Conclusion

The paucity of publicly available environmental monitoring and ecotoxicological data is an obstacle to assessing the risk to the environment posed by pharmaceuticals. Significant data gaps exist, and attempts to use *in-silico* methods to bridge the gaps may contribute to inaccuracy. The worst-case-scenario exposure model utilized by the EMA is based on recommended daily dose, and is so conservative that 98% of pharmaceuticals exceed thresholds for the low-data demand Phase 1 ERA screen.

Medicines approved for sale in the EEA exhibit a distribution of therapeutic uses not reflected in the experimentally derived data for exposure or ecotoxicity of pharmaceuticals that are publicly available. Ecotoxicological data are weighted toward pesticides and endocrine disrupting compounds, while monitoring program data emphasize anti-depressants, anti-epileptics, opioids and other pharmaceuticals acting on the nervous system. One concerning group of pharmaceuticals is the 68 compounds which are demonstrably present in EU surface waters, but for which no ecotoxicity data is available and hence the risk they pose to the environment cannot be assessed.

For those pharmaceuticals which have been reported in monitoring programs since 1997, detection rates approach 50%. Where the risk can be reliably assessed using empirically-derived data, nearly a third of detections exceed their respective risk thresholds. A similarly concerning scenario may also be the case for the 1201 pharmaceuticals which are registered for sale in the EEA, are hence likely to be present in EU surface waters, but for which the exposure and ecotoxicity are unknown.

Prioritizing pharmaceuticals for empirical ecotoxicological studies based on environmental detections, while making a contribution to decreasing the data-gap, is a long term and time-consuming process that would indicate a need for *in-silico* method development. However, the reliability of models and their respective data demands are also likely related. Submission of data to central, publicly accessible databases such as EMPODAT and UBA is voluntary. The pharmaceutical data gap may be alleviated somewhat in future with the introduction of mandatory reporting requirements.

## References

Ågerstrand, M., Berg, C., Björlenius, B., Breitholtz, M., Brunström, B., Fick, J., Gunnarsson, L., Larsson, D.J., Sumpter, J.P., Tysklind, M. and Rudén, C., 2015. Improving environmental risk assessment of human pharmaceuticals. Environmental Science & Technology, 49(9), pp.5336–5345.

Alves, N., Neuparth, T., Barros, S. and Santos, M.M., 2021. The anti-lipidemic drug simvastatin modifies epigenetic biomarkers in the amphipod Gammarus locusta. Ecotoxicology and Environmental Safety, 209, pp.111849.

aus der Beek, T., Weber, F.A., Bergmann, A., Hickmann, S., Ebert, I., Hein, A. and Küster, A., 2016. Pharmaceuticals in the environment—Global occurrences and perspectives. Environmental Toxicology and Chemistry, 35(4), pp.823–835.

Backhaus, T., 2014. Medicines, shaken and stirred: a critical review on the ecotoxicology of pharmaceutical mixtures. Philosophical Transactions of the Royal Society B: Biological Sciences, 369(1656), p.20130585.

Backhaus, T., 2016. Environmental risk assessment of pharmaceutical mixtures: demands, gaps, and possible bridges. The AAPS Journal, 18(4), pp.804–813.

Boxall, A.B., Rudd, M.A., Brooks, B.W., Caldwell, D.J., Choi, K., Hickmann, S., Innes, E., Ostapyk, K., Staveley, J.P., Verslycke, T. and Ankley, G.T., 2012. Pharmaceuticals and personal care products in the environment: what are the big questions? Environmental Health Perspectives, 120(9), pp.1221–1229.

Brack, W., Dulio, V., Ågerstrand, M., Allan, I., Altenburger, R., Brinkmann, M., Bunke, D., Burgess, R.M., Cousins, I., Escher, B.I. and Hernández, F.J., 2017. Towards the review of the European Union Water Framework Directive: recommendations for more efficient assessment and management of chemical contamination in European surface water resources. Science of the Total Environment, 576, pp.720–737.

Brack, W., Hollender, J., de Alda, M.L., Müller, C., Schulze, T., Schymanski, E., Slobodnik, J. and Krauss, M., 2019. High-resolution mass spectrometry to complement monitoring and track emerging chemicals and pollution trends in European water resources. Environmental Sciences Europe, 31(1), pp.1–6.

Brennan, S.J., Brougham, C.A., Roche, J.J. and Fogarty, A.M., 2006. Multi-generational effects of four selected environmental oestrogens on Daphnia magna. Chemosphere, 64(1), pp.49–55.

Burns, E.E., Thomas-Oates, J., Kolpin, D.W., Furlong, E.T. and Boxall, A.B., 2017. Are exposure predictions, used for the prioritization of pharmaceuticals in the environment, fit for purpose? Environmental Toxicology and Chemistry, 36(10), pp.2823–2832.

Burns, E.E., Carter, L.J., Kolpin, D.W., Thomas-Oates, J. and Boxall, A.B., 2018. Temporal and spatial variation in pharmaceutical concentrations in an urban river system. Water Research, 137, pp.72–85.

Caldwell, D.J., Mastrocco, F., Margiotta-Casaluci, L. and Brooks, B.W., 2014. An integrated approach for prioritizing pharmaceuticals found in the environment for risk assessment, monitoring and advanced research. Chemosphere, 115, pp.4–12.

Coquery, M., Morin, A., Becue, A. and Lepot, B., 2005. Priority substances of the European Water Framework Directive: analytical challenges in monitoring water quality. TrAC Trends in Analytical Chemistry, 24(2), pp.117–127.

David, A. and Pancharatna, K., 2009. Effects of acetaminophen (paracetamol) in the embryonic development of zebrafish, Danio rerio. Journal of Applied Toxicology, 29(7), pp.597–602.

De Orte, M.R., Carballeira, C., Viana, I.G. and Carballeira, A., 2013. Assessing the toxicity of chemical compounds associated with marine land-based fish farms: The use of mini-scale microalgal toxicity tests. Chemistry and Ecology, 29(6), pp.554–563.

Dietrich, S., Ploessl, F., Bracher, F. and Laforsch, C., 2010. Single and combined toxicity of pharmaceuticals at environmentally relevant concentrations in Daphnia magna–A multigenerational study. Chemosphere, 79(1), pp.60–66.

Dong, Z., Senn, D.B., Moran, R.E. and Shine, J.P., 2013. Prioritizing environmental risk of prescription pharmaceuticals. Regulatory Toxicology and Pharmacology, 65(1), pp.60–67.

Dortland, R.J., 1980. Toxicological evaluation of parathion and azinphosmethyl in freshwater model ecosystems. Available from ProQuest One Academic. (2565112475). Retrieved from https://www.proquest.com/dissertations-theses/toxicological-evaluation-parathion-azinphosmethyl/docview/2565112475/se-2

Dulio, Valeria, C. Peter, Fabrizio Botta, I. Ipolyi, H. Ruedel, and Jaroslav Slobodnik. “The NORMAN network’s special view on prioritisation of biocides as emerging contaminants.” In 26. SETAC Europe annual meeting. SETAC, 2016.

Dulio, V., van Bavel, B., Brorström-Lundén, E., Harmsen, J., Hollender, J., Schlabach, M., Slobodnik, J., Thomas, K. and Koschorreck, J., 2018. Emerging pollutants in the EU: 10 years of NORMAN in support of environmental policies and regulations. Environmental Sciences Europe, 30(1), pp.1–13.

Dusi, E., Rybicki, M. and Jungmann, D., 2019. The Database” Pharmaceuticals in the Environment”-Update and New Analysis. Umweltbundesamt.

Escher, B.I. and Fenner, K., 2011. Recent advances in environmental risk assessment of transformation products. Environmental Science & Technology, 45(9), pp.3835–3847.

Escher, B. I., Baumgartner, R., Koller, M., Treyer, K., Lienert, J., & McArdell, C. S., 2011. Environmental toxicology and risk assessment of pharmaceuticals from hospital wastewater. Water research, 45(1), 75–92.

EMA/Committee for Medicinal Products for Human Use, 2018. Guideline on the environmental risk assessment of medicinal products for human use - Draft. European Medicines Agency.

European Medecines Agency (EMA), 2023, Article 57 Database, available from: https://www.ema.europa.eu/en/human-regulatory/post-authorisation/data-medicines-iso-idmp-standards/public-data-article-57-database

EC, 2009. Commission Directive 2009/90/EC of 31 July 2009 laying down, pursuant to Directive 2000/60/EC of the European Parliament and of the Council, technical specifications for chemical analysis and monitoring of water status. Official Journal of the European Union L, 201, pp.36.

EC, The Green Deal, Communication COM/2019/640 final. 2019a.

EC, European Union Strategic Approach to Pharmaceuticals in the Environment, Communication COM/2019/128 final. 2019b

EC, Chemicals Strategy for Sustainability Towards a Toxic-Free Environment, Communication COM/2020/667 final. 2020.

EMA/Committee for Medicinal Products for Human Use, 2006. Guideline on the environmental risk assessment of medicinal products for human use. European Medicines Agency.

Felis, E., Kalka, J., Sochacki, A., Kowalska, K., Bajkacz, S., Harnisz, M. and Korzeniewska, E., 2020. Antimicrobial pharmaceuticals in the aquatic environment-occurrence and environmental implications. European Journal of Pharmacology, 866, pp.172813.

Flippin, J.L., Huggett, D. and Foran, C.M., 2007. Changes in the timing of reproduction following chronic exposure to ibuprofen in Japanese medaka, Oryzias latipes. Aquatic Toxicology, 81(1), pp.73–78.

Garric, J., Vollat, B., Duis, K., Péry, A., Junker, T., Ramil, M., Fink, G. and Ternes, T.A., 2007. Effects of the parasiticide ivermectin on the cladoceran Daphnia magna and the green alga Pseudokirchneriella subcapitata. Chemosphere, 69(6), pp.903–910.

Gaston, L., Lapworth, D.J., Stuart, M. and Arnscheidt, J., 2019. Prioritization approaches for substances of emerging concern in groundwater: a critical review. Environmental Science & Technology, 53(11), pp.6107–6122.

González-Pleiter, M., Gonzalo, S., Rodea-Palomares, I., Leganés, F., Rosal, R., Boltes, K., Marco, E. and Fernández-Piñas, F., 2013. Toxicity of five antibiotics and their mixtures towards photosynthetic aquatic organisms: implications for environmental risk assessment. Water Research, 47(6), pp.2050–2064.

Gustavsson, M., Kreuger, J., Bundschuh, M. and Backhaus, T., 2017. Pesticide mixtures in the Swedish streams: environmental risks, contributions of individual compounds and consequences of single-substance oriented risk mitigation. Science of the Total Environment, 598, pp.973–983.

Helsel, D.R., 2010. Summing nondetects: Incorporating low-level contaminants in risk assessment. Integrated Environmental Assessment and Management, 6(3), pp.361–366.

Hollender, J., Zimmermann, S.G., Koepke, S., Krauss, M., McArdell, C.S., Ort, C., Singer, H., Von Gunten, U. and Siegrist, H., 2009. Elimination of organic micropollutants in a municipal wastewater treatment plant upgraded with a full-scale post-ozonation followed by sand filtration. Environmental Science & Technology, 43(20), pp.7862–7869.

Huggett, D.B., Brooks, B.W., Peterson, B., Foran, C.M. and Schlenk, D., 2002. Toxicity of select beta adrenergic receptor-blocking pharmaceuticals (B-blockers) on aquatic organisms. Archives of Environmental Contamination and Toxicology, 43(2), pp.229–235.

Hughes, S.R., Kay, P. and Brown, L.E., 2013. Global synthesis and critical evaluation of pharmaceutical data sets collected from river systems. Environmental Science & Technology, 47(2), pp.661–677.

Isidori, M., Nardelli, A., Pascarella, L., Rubino, M. and Parrella, A., 2007. Toxic and genotoxic impact of fibrates and their photoproducts on non-target organisms. Environment International, 33(5), pp.635–641.

ISO/TS 13530:2009. Water quality — Guidance on analytical quality control for chemical and physicochemical water analysis. Available from: https://www.iso.org/standard/52910.html

Johnson, D.J., Sanderson, H., Brain, R.A., Wilson, C.J. and Solomon, K.R., 2007. Toxicity and hazard of selective serotonin reuptake inhibitor antidepressants fluoxetine, fluvoxamine, and sertraline to algae. Ecotoxicology and Environmental Safety, 67(1), pp.128–139.

Kar, S., Roy, K., & Leszczynski, J., 2018. Impact of pharmaceuticals on the environment: risk assessment using QSAR modeling approach. Computational Toxicology: Methods and Protocols, 395–443.

Kühn, R., Pattard, M., Pernak, K.D. and Winter, A., 1989. Results of the harmful effects of water pollutants to Daphnia magna in the 21-day reproduction test. Water Research, 23(4), pp.501–510.

Kümmerer, K. ed., 2008. Pharmaceuticals in the environment: sources, fate, effects and risks. Springer Science & Business Media.

Kusk, K.O., Christensen, A.M. and Nyholm, N., 2018. Algal growth inhibition test results of 425 organic chemical substances. Chemosphere, 204, pp.405–412.

Küster A, Alder AC, Escher BI, Duis K, Fenner K, Garric J, Hutchinson TH, Lapen DR, Péry A, Römbke J, Snape J, Ternes T, Topp E, Wehrhan A, Knacker T. Environmental risk assessment of human pharmaceuticals in the European Union: A case study with the β-blocker atenolol. Integrated Environmental Assessment and Management. 2010 Jul;6 Suppl:514–23.

Küster, A., & Adler, N. (2014). Pharmaceuticals in the environment: scientific evidence of risks and its regulation. Philosophical Transactions of the Royal Society B: Biological Sciences, 369(1656), 20130587.

Ladouceur, M.F., 1992. Biodegradation of the organophosphorus insecticide fenitrothion by the alga Chlamydomonas reinhardtii: The role of cytochrome P450 monooxygenase. (Doctoral dissertation, University of Ottawa).

Lange, A., Paull, G.C., Coe, T.S., Katsu, Y., Urushitani, H., Iguchi, T. and Tyler, C.R., 2009. Sexual reprogramming and estrogenic sensitization in wild fish exposed to ethinylestradiol. Environmental Science & Technology, 43(4), pp.1219–1225.

Lister, A.L., 2009. Prostaglandins in the zebrafish ovary: Role, regulation, and modulation by environmental pharmaceuticals (Doctoral dissertation, University of Guelph).

Loos, R., Gawlik, B.M., Locoro, G., Rimaviciute, E., Contini, S. and Bidoglio, G., 2009. EU-wide survey of polar organic persistent pollutants in European river waters. Environmental Pollution, 157(2), pp.561–568.

Loos, R., Carvalho, R., António, D.C., Comero, S., Locoro, G., Tavazzi, S., Paracchini, B., Ghiani, M., Lettieri, T., Blaha, L. and Jarosova, B., 2013. EU-wide monitoring survey on emerging polar organic contaminants in wastewater treatment plant effluents. Water Research, 47(17), pp.6475–6487.

Loos, R., Marinov, D., Sanseverino, I., Napierska, D. and Lettieri, T., 2018. Review of the 1st Watch List under the Water Framework Directive and recommendations for the 2nd Watch List. Luxembourg: Publications Office of the European Union.

Lu, G., Li, Z. and Liu, J., 2013. Effects of selected pharmaceuticals on growth, reproduction and feeding of Daphnia magna. Fresenius Environmental Bulletin, 22(09), pp.2588–2594.

Mansour, F., Al-Hindi, M., Saad, W. and Salam, D., 2016. Environmental risk analysis and prioritization of pharmaceuticals in a developing world context. Science of the Total Environment, 557, pp.31–43.

Mallett, M.J. and Sims, I., 1994. Effects of ammonia on the early life stage of carp (Cyprinus carpio) and roach (Rutilus rutilus). In: R. Muller and R. Lloyd (Eds.), Sublethal and Chronic Effects of Pollutants on Freshwater Fish, Fishing News Books, London 19:211–228

Mitchell, R.J., Myers, A.L., Mabury, S.A., Solomon, K.R. and Sibley, P.K., 2011. Toxicity of fluorotelomer carboxylic acids to the algae Pseudokirchneriella subcapitata and Chlorella vulgaris, and the amphipod Hyalella azteca. Ecotoxicology and Environmental Safety, 74(8), pp.2260–2267.

Overturf, M.D., Overturf, C.L., Baxter, D., Hala, D.N., Constantine, L., Venables, B. and Huggett, D.B., 2012. Early life-stage toxicity of eight pharmaceuticals to the fathead minnow, Pimephales promelas. Archives of Environmental Contamination and Toxicology, 62(3), pp.455–464.

Park, S. and Choi, K., 2008. Hazard assessment of commonly used agricultural antibiotics on aquatic ecosystems. Ecotoxicology, 17(6), pp.526–538.

Perazzolo, C., Morasch, B., Kohn, T., Magnet, A., Thonney, D. and Chèvre, N., 2010. Occurrence and fate of micropollutants in the Vidy Bay of Lake Geneva, Switzerland. Part I: priority list for environmental risk assessment of pharmaceuticals. Environmental Toxicology and Chemistry, 29(8), pp.1649–1657.

Pereira, A., Silva, L., Laranjeiro, C., Lino, C. and Pena, A., 2020a. Selected pharmaceuticals in different aquatic compartments: Part I—Source, fate and occurrence. Molecules, 25(5), p.1026.

Pereira, A., Silva, L., Laranjeiro, C., Lino, C. and Pena, A., 2020b. Selected pharmaceuticals in different aquatic compartments: Part II—Toxicity and environmental risk assessment. Molecules, 25(8), p.1796.

Posthuma, L., van Gils, J., Zijp, M.C., van De Meent, D. and de Zwart, D., 2019. Species sensitivity distributions for use in environmental protection, assessment, and management of aquatic ecosystems for 12 386 chemicals. Environmental Toxicology and Chemistry, 38(4), pp.905–917.

Roos, V., Gunnarsson, L., Fick, J., Larsson, D.G.J. and Rudén, C., 2012. Prioritising pharmaceuticals for environmental risk assessment: towards adequate and feasible first-tier selection. Science of the Total Environment, 421, pp.102–110.

Runnalls, T.J., Margiotta-Casaluci, L., Kugathas, S. and Sumpter, J.P., 2010. Pharmaceuticals in the aquatic environment: steroids and anti-steroids as high priorities for research. Human and Ecological Risk Assessment, 16(6), pp.1318–1338.

Russo, C., Lavorgna, M., Česen, M., Kosjek, T., Heath, E. and Isidori, M., 2018. Evaluation of acute and chronic ecotoxicity of cyclophosphamide, ifosfamide, their metabolites/transformation products and UV treated samples. Environmental Pollution, 233, pp.356–363.

Sanderson, H., & Thomsen, M., 2007. Ecotoxicological quantitative structure–activity relationships for pharmaceuticals. Bulletin of Environmental Contamination and Toxicology, 79, 331–335.

Schreiner, V.C., Szöcs, E., Bhowmik, A.K., Vijver, M.G. and Schäfer, R.B., 2016. Pesticide mixtures in streams of several European countries and the USA. Science of the Total Environment, 573, pp.680–689.

Schwarz, S., Gildemeister, D., Hein, A., Schröder, P., & Bachmann, J. (2021). Environmental fate and effects assessment of human pharmaceuticals: lessons learnt from regulatory data. Environmental Sciences Europe, 33(1), 1–20.

Shioda, T. and Wakabayashi, M., 2000. Evaluation of reproductivity of medaka (Oryzias latipes) exposed to chemicals using a 2-week reproduction test. Water Science and Technology, 42(7-8), pp.53–60.

Spilsbury, F.D., Warne, M.S.J. and Backhaus, T., 2020. Risk assessment of pesticide mixtures in Australian rivers discharging to the Great Barrier Reef. Environmental Science & Technology, 54(22), pp.14361–14371.

Steinkey, D., Lari, E., Woodman, S.G., Luong, K.H., Wong, C.S. and Pyle, G.G., 2018. Effects of gemfibrozil on the growth, reproduction, and energy stores of Daphnia magna in the presence of varying food concentrations. Chemosphere, 192, pp.75–80.

Szöcs E, Stirling T, Scott ER, et al. (2020) webchem: An R Package to Retrieve Chemical Information from the Journal of Statistical Software 93:. https://doi.org/10.18637/jss.v093.i13

UBA (2022), Database – Pharmaceuticals in the Environment. Available from: https://www.umweltbundesamt.de/en/database-pharmaceuticals-in-the-environment-0; date accessed: 8th September 2022.

U.S. EPA, 1992. Pesticide Ecotoxicity Database (Formerly: Environmental Effects Database (EEDB)), Environmental Fate and Effects Division, U.S.EPA, Washington, D.C.:

U.S. EPA, ECOTOX User Guide: ECOTOXicology Knowledgebase System. Version 5.3. http://www.epa.gov/ecotox/ 2022.

von der Ohe, P.C., Dulio, V., Slobodnik, J., De Deckere, E., Kühne, R., Ebert, R.U., Ginebreda, A., De Cooman, W., Schüürmann, G. and Brack, W., 2011. A new risk assessment approach for the prioritization of 500 classical and emerging organic microcontaminants as potential river basin specific pollutants under the European Water Framework Directive. Science of the Total Environment, 409(11), pp.2064–2077.

Yang, L.H., Ying, G.G., Su, H.C., Stauber, J.L., Adams, M.S. and Binet, M.T., 2008. Growth-inhibiting effects of 12 antibacterial agents and their mixtures on the freshwater microalga Pseudokirchneriella subcapitata. Environmental Toxicology and Chemistry, 27(5), pp.1201–1208.

