## Supplemental Figures S1, S2 and Tables S1, S2 and S3 for "Defining the data gap: what do we know about environmental exposure, hazards and risks of pharmaceuticals in the European aquatic environment?"

### Supplementary Information

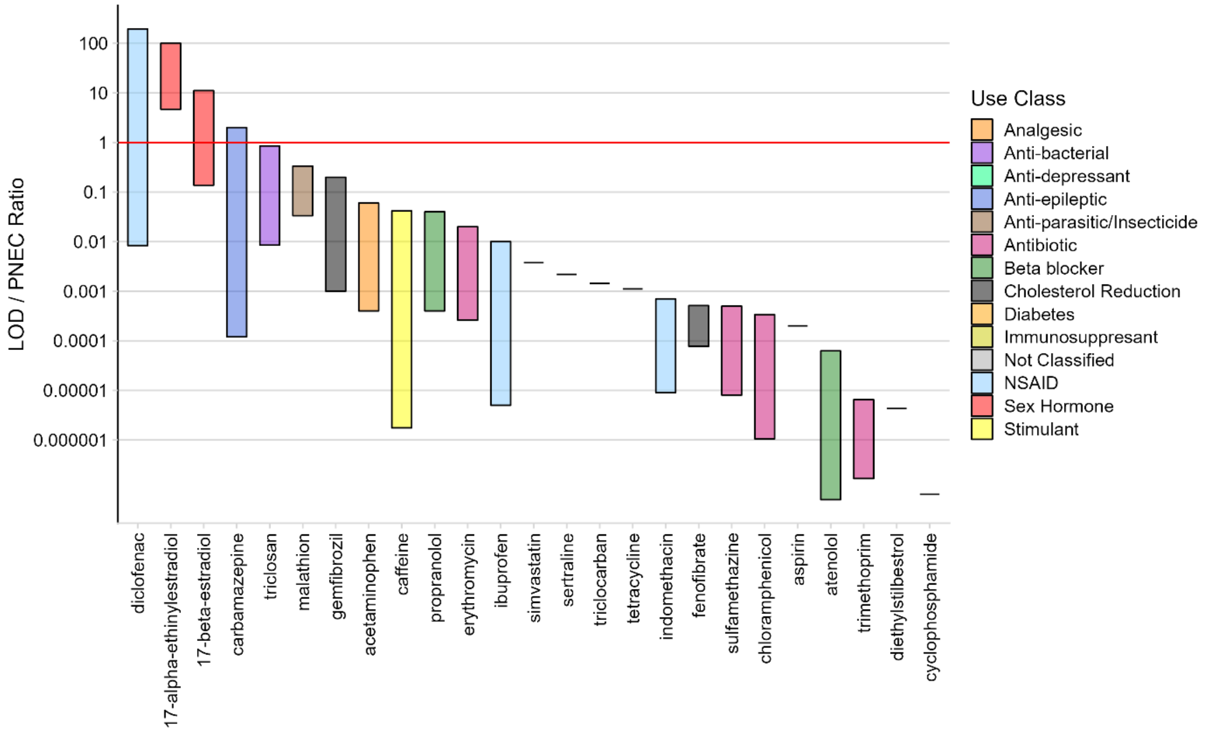


Figure S1 LOD/PNEC ratios of 25 Active Pharmaceutical Ingredients

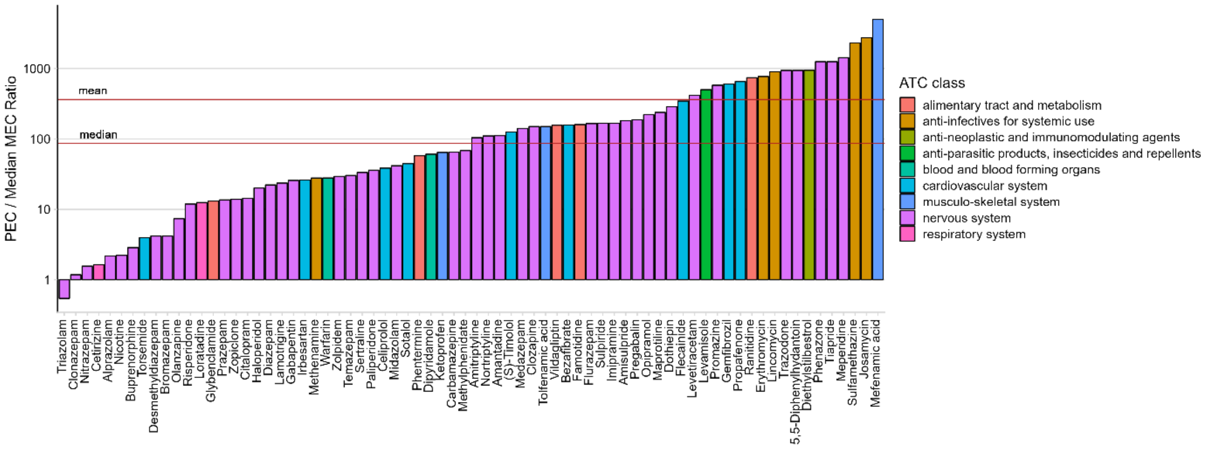


Figure S2 MEC/PEC ratios of 72 Active Pharmaceutical Ingredients

Table S1: Ecotoxicological PNEC data used for risk assessment

| Name | CAS Number | Test Species | Common Name | Endpoint | Measurement | PNEC* (ug/L) | Reference |
|---|---|---|---|---|---|---|---|
| 2-Phenylphenol | 90-43-7 | *Raphidocelis subcapitata* | Green Algae | NOEL | Abundance | 43.2 | EPA, 1992 |
| Ammonia | 7664-41-7 | *Rutilus rutilus* | Roach | NOEC | Mortality | 55.0 | Mallet & Sims, 1994 |
| Aspirin | 50-78-2 | *Danio rerio* | Zebra Danio | NOEC | Progeny counts/numbers | 10.0 | Lister, 2009 |
| Atenolol | 29122-68-7 | *Daphnia magna* | Water Flea | NOEC | Reproduction, general | 320 | Kuster *et al.*, 2010 |
| beta-Estradiol | 50-28-2 | *Oryzias latipes* | Japanese Medaka | NOEC | Hatch | $3 \times 10^{-4}$ | Shioda & Wakabayashi, 2000 |
| Bezafibrate | 41859-67-0 | *Ceriodaphnia dubia* | Water Flea | NOEC | Population growth rate | 2.300 | Isidori *et al.*, 2007 |
| Bronopol | 52-51-7 | *Anabaena flosaquae* | Blue-Green Algae | NOEL | Abundance | 1.20 | EPA, 1992 |
| Caffeine | 58-08-2 | *Daphnia magna* | Water Flea | NOEC | Length | 12.0 | Lu *et al.*, 2013 |
| Carbamazepine | 298-46-4 | *Daphnia magna* | Water Flea | NOEC | Progeny counts/numbers | 0.050 | Dietrich *et al.*, 2010 |
| Chloramphenicol | 56-75-7 | *Raphidocelis subcapitata* | Green Algae | EC10 | Population growth rate | 19.0 | Kusk *et al.,* 2018 |
| Chlorocresol | 59-50-7 | *Daphnia magna* | Water Flea | NOEC | Progeny counts/numbers | 130 | Kuhn *et al.*, 1989 |
| Cyclophosphamide | 50-18-0 | *Raphidocelis subcapitata* | Green Algae | NOEC | Abundance | 1250 | Russo *et al.*, 2018 |
| Diclofenac | 15307-86-5 | *Daphnia magna* | Water Flea | NOEC | Progeny counts/numbers | 0.036 | Dietrich *et al.*, 2010 |
| Diethylstilbestrol | 56-53-1 | *Daphnia magna* | Water Flea | NOEC | Progeny counts/numbers | 10.0 | Brennan *et al.*, 2006 |
| Erythromycin | 114-07-8 | *Anabaena sp.* | Blue-Green Algae | EC10 | Population growth rate | 0.500 | Gonzalez-Pleiter *et al.*, 2013 |
| Ethynyl Estradiol | 57-63-6 | *Rutilus rutilus* | Roach | NOEC | Length | $3 \times 10^{-5}$ | Lange *et al.*, 2009 |
| Fenofibrate | 49562-28-9 | *Pimephales promelas* | Fathead Minnow | NOEC | Survival | 16.9 | Overturf *et al.*, 2012 |
| Formaldehyde | 50-00-0 | *Isochrysis galbana* | Haptophyte | NOEC | Abundance | 0.500 | De Orte *et al.*, 2013 |
| Gemfibrozil | 25812-30-0 | *Daphnia magna* | Water Flea | NOEC | Progeny counts/numbers | 0.050 | Steinkey *et al.*, 2018 |
| Ibuprofen | 15687-27-1 | *Oryzias latipes* | Japanese Medaka | NOEC | Progeny counts/numbers | 10.0 | Flippin, *et al.*, 2007 |
| Indometacin | 53-86-1 | *Danio rerio* | Zebra Danio | NOEC | Progeny counts/numbers | 10.0 | Lister, 2009 |
| Isopropanol | 67-63-0 | *Raphidocelis subcapitata* | Green Algae | NOEC | Abundance | 1000 | Mitchell *et al.,* 2011 |
| Ivermectin | 70288-86-7 | *Daphnia magna* | Water Flea | NOEC | Progeny counts/numbers | $3 \times 10^{-8}$ | Garric *et al.*, 2007 |
| Malathion | 121-75-5 | *Daphnia magna* | Water Flea | NOEC | Reproduction, general | 0.030 | Dortland, 1980 |
| Niclosamide | 50-65-7 | *Daphnia magna* | Water Flea | NOEC | Progeny counts/numbers | 2.00 | Kuhn *et al.*, 1989 |
| Paracetamol | 103-90-2 | *Danio rerio* | Zebra Danio | NOEC | Hatch | 0.100 | David & Pancharatna, 2009 |
| Permethrin | 52645-53-1 | *Daphnia magna* | Water Flea | NOEC | Hatch | 0.004 | EPA, 1992 |

| | | | | | | | |
|---|---|---|---|---|---|---|---|
| Phenobarbital | 50-06-6 | *Chlamydomonas reinhardtii* | Green Algae | NOEC | Biomass | 2320 | Ladouceur (1992) |
| Propranolol | 525-66-6 | *Hyalella azteca* | Scud | NOEC | Progeny counts/numbers | 0.100 | Huggett *et al.*, 2002 |
| Sertraline | 79617-96-2 | *Raphidocelis subcapitata* | Green Algae | IC10 | Abundance | 0.457 | Johnson *et al.*, 2007 |
| Simvastatin | 79902-63-9 | *Gammarus locusta* | Scud | NOEC | Survival | 0.800 | Alves *et al.*, 2021 |
| Sulfamethazine | 57-68-1 | *Raphidocelis subcapitata* | Green Algae | NOEC | Biomass | 100 | Yang *et al.*, 2008 |
| Tetracycline | 60-54-8 | *Raphidocelis subcapitata* | Green Algae | EC10 | Population growth rate | 3.20 | Gonzalez-Pleiter *et al.*, 2013 |
| Thiram | 137-26-8 | *Pimephales promelas* | Fathead Minnow | NOEC | Progeny counts/numbers | 0.110 | EPA, 1992 |
| Trichlocarban | 101-20-2 | *Daphnia magna* | Water Flea | NOEC | Progeny counts/numbers | 0.025 | EPA, 1992 |
| Triclosan | 3380-34-5 | *Raphidocelis subcapitata* | Green Algae | NOEC | Biomass | 0.020 | Yang *et al.*, 2008 |
| Trimethoprim | 738-70-5 | *Daphnia magna* | Water Flea | NOEC | Progeny counts/numbers | 600 | Park & Choi, 2008 |

*PNECs were calculated as chronic NOEC or EC10 for the most sensitive species, with an assessment factor of 10 (EMA, 2018).

Table S2: APIs detected in European surface waters (1997 – 2020) lacking ecotoxicological data

| Name of Analyte | CAS number | Number Monitoring Records | Number Detections | Detection Rate (%) | Median MEC (µg/L) | ATC Group1 | ATC Group3 |
|---|---|---|---|---|---|---|---|
| Acetazolamide | 59-66-5 | 343 | 54 | 15.74 | 0.003 | genito urinary system and sex hormones | anti-infectives and anti-septics, excl. corticosteroids |
| Albuterol | 18559-94-9 | 308 | 62 | 20.13 | 0.0005 | respiratory system | adrenergics, inhalants |
| Alfentanil | 71195-58-9 | 68 | 36 | 52.94 | 0.0008 | nervous system | anesthetics, general |
| Alprazolam | 28981-97-7 | 914 | 218 | 23.85 | 0.002 | nervous system | anxiolytics |
| Amisulpride | 71675-85-9 | 337 | 193 | 57.27 | 0.010 | nervous system | anti-psychotics |
| Azidothymidine | 30516-87-1 | 117 | 1 | 0.85 | 0.002 | anti-infectives for systemic use | direct acting anti-virals |
| Bromazepam | 1812-30-2 | 499 | 62 | 12.42 | 0.012 | nervous system | anxiolytics |
| Buprenorphine | 52485-79-7 | 181 | 64 | 35.36 | 0.012 | nervous system | drugs used in addictive disorders |
| Candesartan | 139481-59-7 | 483 | 294 | 60.87 | 0.031 | cardiovascular system | angiotensin ii receptor blockers (arbs), combinations |
| Celiprolol | 56980-93-9 | 436 | 397 | 91.06 | 0.026 | cardiovascular system | beta blocking agents |
| Cetirizine | 83881-51-0 | 680 | 403 | 59.26 | 0.030 | respiratory system | anti-histamines for systemic use |
| Cilastatin | 82009-34-5 | 119 | 2 | 1.68 | 0.058 | anti-infectives for systemic use | other beta-lactam anti-bacterials |
| Clonazepam | 1622-61-3 | 146 | 31 | 21.23 | 0.022 | nervous system | anti-epileptics |
| Codeine | 76-57-3 | 509 | 261 | 51.28 | 0.012 | nervous system | opioids |
| Crotamiton | 483-63-6 | 111 | 4 | 3.60 | 0.007 | various / uncategorized | uncategorized |
| Dapsone | 80-08-0 | 310 | 1 | 0.32 | 0.50 | dermatologicals | anti-acne preparations for topical use |
| Nordazepem | 1088-11-5 | 191 | 130 | 68.06 | 0.017 | nervous system | anxiolytics |
| Dihydrocodeine | 125-28-0 | 130 | 124 | 95.38 | 0.39 | nervous system | opioids |
| Dipyridamole | 58-32-2 | 184 | 1 | 0.54 | 0.033 | blood and blood forming organs | anti-thrombotic agents |
| Dothiepin | 113-53-1 | 65 | 64 | 98.46 | 0.003 | nervous system | anti-depressants |
| Esmolol | 81147-92-4 | 116 | 1 | 0.86 | 0.081 | cardiovascular system | beta blocking agents |
| Fentanyl | 437-38-7 | 65 | 18 | 27.69 | 0.0008 | uncategorized | uncategorized |
| Flecainide | 54143-55-4 | 3 | 1 | 33.33 | 0.003 | cardiovascular system | anti-arrhythmics, class i and iii |
| Flunitrazepam | 1622-62-4 | 65 | 3 | 4.62 | 0.001 | nervous system | hypnotics and sedatives |

| | | | | | | | |
|---|---|---|---|---|---|---|---|
| Flurazepam | 17617-23-1 | 81 | 1 | 1.23 | 0.0009 | nervous system | hypnotics and sedatives |
| Gestodene | 60282-87-3 | 22 | 1 | 4.55 | 0.004 | genito urinary system and sex hormones | hormonal contraceptives for systemic use |
| Heroin | 561-27-3 | 155 | 1 | 0.65 | 0.002 | nervous system | drugs used in addictive disorders |
| Hydromorphone | 466-99-9 | 107 | 45 | 42.06 | 0.001 | nervous system | opioids |
| Iomeprol | 78649-41-9 | 2944 | 2412 | 81.93 | 0.23 | various / uncategorized | x-ray contrast media, iodinated |
| Iopamidol | 60166-93-0 | 2240 | 1448 | 64.64 | 0.13 | various / uncategorized | x-ray contrast media, iodinated |
| Josamycin | 16846-24-5 | 25 | 4 | 16.00 | 0.004 | anti-infectives for systemic use | macrolides, lincosamides and streptogramins |
| Ketamine | 6740-88-1 | 181 | 27 | 14.92 | 0.001 | uncategorized | uncategorized |
| Lamotrigine | 84057-84-1 | 1116 | 1040 | 93.19 | 0.060 | nervous system | anti-epileptics |
| Levetiracetam | 102767-28-2 | 3 | 1 | 33.33 | 0.018 | nervous system | anti-epileptics |
| Lopinavir | 192725-17-0 | 55 | 1 | 1.82 | 0.077 | anti-infectives for systemic use | direct acting anti-virals |
| Lorazepam | 846-49-1 | 1331 | 294 | 22.09 | 0.011 | nervous system | anxiolytics |
| Maprotiline | 10262-69-8 | 65 | 51 | 78.46 | 0.002 | nervous system | anti-depressants |
| Medazepam | 2898-12-6 | 367 | 48 | 13.08 | 0.0006 | nervous system | anxiolytics |
| Meperidine | 57-42-1 | 245 | 57 | 23.27 | 0.001 | nervous system | opioids |
| Mestranol | 72-33-3 | 568 | 84 | 14.79 | 0.10 | uncategorized | uncategorized |
| Ritalin | 113-45-1 | 65 | 46 | 70.77 | 0.002 | nervous system | psychostimulants, agents for adhd and nootropics |
| Midazolam | 59467-70-8 | 65 | 5 | 7.69 | 0.002 | nervous system | hypnotics and sedatives |
| Niflumic acid | 4394-00-7 | 116 | 5 | 4.31 | 0.0002 | musculo-skeletal system | topical products for joint and muscular pain |
| Nitrazepam | 146-22-5 | 65 | 44 | 67.69 | 0.015 | nervous system | hypnotics and sedatives |
| Olanzapine | 132539-06-1 | 184 | 57 | 30.98 | 0.006 | nervous system | anti-psychotics |
| Opipramol | 315-72-0 | 181 | 24 | 13.26 | 0.004 | nervous system | anti-depressants |
| Paliperidone | 144598-75-4 | 163 | 26 | 15.95 | 0.0008 | nervous system | anti-psychotics |
| Pentazocine | 359-83-1 | 78 | 7 | 8.97 | 0.0004 | uncategorized | uncategorized |
| Phenazepam | 51753-57-2 | 181 | 26 | 14.36 | 0.011 | uncategorized | uncategorized |
| Phenazone | 60-80-0 | 2103 | 804 | 38.23 | 0.011 | nervous system | other analgesics and anti-pyretics |
| Phentermine | 122-09-8 | 139 | 25 | 17.99 | 0.001 | alimentary tract and metabolism | anti-obesity preparations, excl. diet products |
| Prazepam | 2955-38-6 | 249 | 72 | 28.92 | 0.013 | nervous system | anxiolytics |

| | | | | | | | |
|---|---|---|---|---|---|---|---|
| Pregabalin | 148553-50-8 | 181 | 8 | 4.42 | 0.008 | nervous system | anti-epileptics |
| Promazine | 58-40-2 | 68 | 32 | 47.06 | 0.002 | nervous system | anti-psychotics |
| Promethazine | 60-87-7 | 65 | 29 | 44.62 | 0.007 | respiratory system | anti-histamines for systemic use |
| Propafenone | 54063-53-5 | 68 | 31 | 45.59 | 0.002 | cardiovascular system | anti-arrhythmics, class i and iii |
| Propyphenazone | 479-92-5 | 11983 | 3657 | 30.52 | 0.010 | nervous system | other analgesics and anti-pyretics |
| Risperidone | 106266-06-2 | 184 | 29 | 15.76 | 0.002 | nervous system | anti-psychotics |
| Sitagliptin | 486460-32-6 | 907 | 876 | 96.58 | 0.11 | alimentary tract and metabolism | blood glucose lowering drugs, excl. insulins |
| Tiapride | 51012-32-9 | 3 | 2 | 66.67 | 0.002 | nervous system | anti-psychotics |
| Timolol | 26839-75-8 | 28 | 6 | 21.43 | 0.0008 | cardiovascular system | beta blocking agents |
| Torsemide | 56211-40-6 | 653 | 504 | 77.18 | 0.019 | cardiovascular system | high-ceiling diuretics |
| Trazodone | 19794-93-5 | 65 | 44 | 67.69 | 0.002 | nervous system | anti-depressants |
| Triazolam | 28911-01-5 | 68 | 37 | 54.41 | 0.002 | nervous system | hypnotics and sedatives |
| Trimipramine | 739-71-9 | 68 | 63 | 92.65 | 0.15 | uncategorized | uncategorized |
| Vildagliptin | 274901-16-5 | 116 | 3 | 2.59 | 0.003 | alimentary tract and metabolism | blood glucose lowering drugs, excl. insulins |
| Zolpidem | 82626-48-0 | 470 | 81 | 17.23 | 0.002 | nervous system | hypnotics and sedatives |
| Zopiclone | 43200-80-2 | 184 | 41 | 22.28 | 0.003 | nervous system | hypnotics and sedatives |

Table S3: APIs prioritized for inclusion in monitoring programs

| Name | CAS number | DDD (mg/day) | PEC (µg/L) | Log Kow | Hormone | Antibiotic | Anti-mycotic | Insectide/ repellent | ATC Group 3 |
|---|---|---|---|---|---|---|---|---|---|
| Amphotericin B | 1397-89-3 | 200 | 1.00 | | × | ✓ | × | × | anti-infectives and anti-septics, excl. corticosteroids |
| Anidulafungin | 166663-25-8 | 100 | 0.50 | | × | × | ✓ | × | anti-mycotics for systemic use |
| Atovaquone | 95233-18-4 | 2250 | 11.25 | 5.80 | × | × | × | ✓ | agents against amoebiasis and other protozoal |
| Aztreonam | 78110-38-0 | 4000 | 20.00 | 0.30 | × | ✓ | × | × | other beta-lactam anti-bacterials |
| Bedaquiline | 843663-66-1 | 86 | 0.43 | | × | ✓ | ✓ | × | drugs for treatment of tuberculosis |
| Capreomycin | 11003-38-6 | 1000 | 5.00 | | × | ✓ | × | × | drugs for treatment of tuberculosis |
| Caspofungin | 162808-62-0 | 50 | 0.25 | | × | × | ✓ | × | anti-mycotics for systemic use |
| Cefaclor | 53994-73-3 | 1000 | 5.00 | -1.80 | × | ✓ | × | × | other beta-lactam anti-bacterials |
| Cefalotin | 153-61-7 | 4000 | 20.00 | | × | ✓ | × | × | other beta-lactam anti-bacterials |
| Cefamandole | 34444-01-4 | 6000 | 30.00 | 0.50 | × | ✓ | × | × | other beta-lactam anti-bacterials |
| Cefatrizine | 51627-14-6 | 1000 | 5.00 | | × | ✓ | × | × | other beta-lactam anti-bacterials |
| Cefepime | 88040-23-7 | 4000 | 20.00 | -0.10 | × | ✓ | × | × | other beta-lactam anti-bacterials |
| Cefmetazole | 56796-20-4 | 4000 | 20.00 | | × | ✓ | × | × | other beta-lactam anti-bacterials |
| Cefodizime | 69739-16-8 | 2000 | 10.00 | 0.20 | × | ✓ | × | × | other beta-lactam anti-bacterials |
| Cefoperazone | 62893-19-0 | 4000 | 20.00 | | × | ✓ | × | × | other beta-lactam anti-bacterials |
| Cefprozil | 92665-29-7 | 1000 | 5.00 | | × | ✓ | × | × | other beta-lactam anti-bacterials |
| Cefradine | 38821-53-3 | 2000 | 10.00 | | × | ✓ | × | × | other beta-lactam anti-bacterials |
| Ceftaroline Fosamil | 229016-73-3 | 1200 | 6.00 | | × | ✓ | × | × | other beta-lactam anti-bacterials |
| Ceftibuten | 97519-39-6 | 400 | 2.00 | | × | ✓ | × | × | other beta-lactam anti-bacterials |
| Ceftobiprole Medocaril | 376653-43-9 | 1500 | 7.50 | | × | ✓ | × | × | other beta-lactam anti-bacterials |
| Clofoctol | 37693-01-9 | 1500 | 7.50 | 8.10 | × | ✓ | × | × | other anti-bacterials |
| Clomifene Citrate | 911-45-5 | 9 | 0.05 | | ✓ | × | × | × | gonadotropins and other ovulation stimulants |
| Danazol | 17230-88-5 | 600 | 3.00 | 4.08 | ✓ | × | × | × | other sex hormones and genital system modulators |
| Daptomycin | 103060-53-3 | 280 | 1.40 | -5.10 | × | ✓ | × | × | other anti-bacterials |

| | | | | | | | | | |
|---|---|---|---|---|---|---|---|---|---|
| Delamanid | 681492-22-8 | 200 | 1.00 | | × | × | ✓ | × | drugs for treatment of tuberculosis |
| Demeclocycline | 127-33-3 | 600 | 3.00 | | × | ✓ | × | × | tetracyclines |
| Diethylcarbamazine | 90-89-1 | 400 | 2.00 | | × | × | × | ✓ | anti-nematodal agents |
| Ertapenem | 153832-46-3 | 1000 | 5.00 | 0.30 | × | ✓ | × | × | other beta-lactam anti-bacterials |
| Ethambutol | 74-55-5 | 1200 | 6.00 | 0.40 | × | ✓ | × | × | drugs for treatment of tuberculosis |
| Ethionamide | 536-33-4 | 750 | 3.75 | | × | ✓ | × | × | drugs for treatment of tuberculosis |
| Flubendazole | 31430-15-6 | 200 | 1.00 | 2.90 | × | × | × | ✓ | anti-nematodal agents |
| Fosfomycin | 23155-02-4 | 8000 | 40.00 | -1.60 | × | ✓ | × | × | other anti-bacterials |
| Furazidine | 1672-88-4 | 300 | 1.50 | 0.50 | × | ✓ | × | × | other anti-bacterials |
| Griseofulvin | 126-07-8 | 500 | 2.50 | 2.18 | × | × | ✓ | × | anti-fungals for systemic use |
| Itraconazole | 84625-61-6 | 200 | 1.00 | 5.70 | × | × | ✓ | × | anti-mycotics for systemic use |
| Lymecycline | 992-21-2 | 600 | 3.00 | | × | ✓ | × | × | tetracyclines |
| Meglumine Antimonate | 133-51-7 | 850 | 4.25 | | × | ✓ | × | ✓ | agents against leishmaniasis and trypanosomiasis |
| Melarsoprol | 494-79-1 | 60 | 0.30 | | × | ✓ | × | ✓ | agents against leishmaniasis and trypanosomiasis |
| Meropenem | 96036-03-2 | 3000 | 15.00 | -0.60 | × | ✓ | × | × | other beta-lactam anti-bacterials |
| Mesterolone | 1424-00-6 | 50 | 0.25 | 4.10 | ✓ | × | × | × | androgens |
| Metacycline | 914-00-1 | 600 | 3.00 | | × | ✓ | × | × | tetracyclines |
| Micafungin | 235114-32-6 | 100 | 0.50 | | × | × | ✓ | × | anti-mycotics for systemic use |
| Mifepristone | 84371-65-3 | 200 | 1.00 | 5.40 | ✓ | × | × | × | other sex hormones and genital system modulators |
| Minocycline | 10118-90-8 | 200 | 1.00 | | × | ✓ | × | × | tetracyclines |
| Natamycin | 7681-93-8 | 25 | 0.13 | | × | ✓ | × | × | anti-infectives and anti-septics, excl. corticosteroids |
| Nifuratel | 4936-47-4 | 600 | 3.00 | 0.70 | × | ✓ | × | × | anti-infectives and anti-septics, excl. corticosteroids |
| Nitrofurantoin | 67-20-9 | 200 | 1.00 | -0.50 | × | ✓ | × | × | other anti-bacterials |
| Paromomycin | 7542-37-2 | 3000 | 15.00 | -8.70 | × | ✓ | × | × | intestinal anti-infectives |
| Pentamidine | 100-33-4 | 280 | 1.40 | | × | × | × | ✓ | agents against leishmaniasis and trypanosomiasis |
| Phenolphthalein | 77-09-8 | 200 | 1.00 | | ✓ | × | × | × | drugs for constipation |
| Posaconazole | 171228-49-2 | 300 | 1.50 | 4.60 | × | × | ✓ | × | anti-mycotics for systemic use |

| | | | | | | | | | |
|---|---|---|---|---|---|---|---|---|---|
| Protionamide | 14222-60-7 | 750 | 3.75 | 1.50 | × | ✓ | × | × | drugs for treatment of tuberculosis |
| Pyrazinamide | 98-96-4 | 1500 | 7.50 | -0.60 | × | ✓ | × | × | drugs for treatment of tuberculosis |
| Pyrvinium | 7187-62-4 | 350 | 1.75 | | × | × | × | ✓ | anti-nematodal agents |
| Rifabutin | 72559-06-9 | 150 | 0.75 | | × | ✓ | × | × | drugs for treatment of tuberculosis |
| Secnidazole | 3366-95-8 | 2000 | 10.00 | 0.22 | × | × | × | ✓ | agents against amoebiasis and other protozoal |
| Spectinomycin | 1695-77-8 | 3000 | 15.00 | | × | ✓ | × | × | other anti-bacterials |
| Spironolactone | 52-01-7 | 75 | 0.38 | | ✓ | × | × | × | aldosterone antagonists and potassium-sparing agents |
| Sulbactam | 68373-14-8 | 1000 | 5.00 | | × | ✓ | × | × | beta-lactam anti-bacterials, penicillins |
| Suramin Sodium | 145-63-1 | 270 | 1.35 | | × | × | × | ✓ | agents against leishmaniasis and trypanosomiasis |
| Teicoplanin | 61036-62-2 | 400 | 2.00 | 0.50 | × | ✓ | × | × | other anti-bacterials |
| Testosterone | 58-22-0 | 120 | 0.60 | 3.32 | ✓ | × | × | × | androgens |
| Tibolone | 5630-53-5 | 2.5 | 0.01 | 2.40 | ✓ | × | × | × | estrogens |
| Ulipristal Acetate | 159811-51-5 | 30 | 0.15 | | ✓ | × | × | × | hormonal contraceptives for systemic use |
| Voriconazole | 137234-62-9 | 400 | 2.00 | 1.00 | × | × | ✓ | × | anti-mycotics for systemic use |
